# Gene duplication accelerates the pace of protein gain and loss from plant organelles

**DOI:** 10.1101/415125

**Authors:** Rona Costello, David M. Emms, Steven Kelly

## Abstract

A hallmark of eukaryotic cells is the compartmentalisation of intracellular processes into specialised membrane-bound compartments known as organelles. Plant cells contain several such organelles including the nucleus, chloroplast, mitochondrion, peroxisome, golgi, endoplasmic reticulum and vacuole. Organelle biogenesis and function is dependent on the concerted action of numerous nuclear-encoded proteins which must be imported from the cytosol (or endoplasmic reticulum) where they are made. Using phylogenomic approaches coupled to ancestral state estimation we show that the rate of change in plant organellar proteome content is proportional to the rate of molecular sequence evolution such that the proteomes of chloroplasts and mitochondria lose or gain ~3.2 proteins per million years. We show that these changes in protein targeting have predominantly occurred in genes with regulatory rather than metabolic functions, and thus altered regulatory capacity rather than metabolic function has been the major theme of plant organellar evolution. Finally we show gain and loss of protein targeting occurs at a higher rate following gene duplication events, revealing that gene and genome duplication are a key facilitator of organelle evolution.

## Main text

While the chloroplast and mitochondrion contain DNA that encodes a number of organellar intrinsic proteins, the vast majority of chloroplast and mitochondrial proteins are encoded in the nucleus ^1^. Moreover, the proteome of the secretory organelles, peroxisome and vacuole is entirely encoded in the nuclear genome. Nuclear-encoded organellar proteins are translocated to and across the organelle membrane by means of a short, often cleavable, targeting signal located within the amino acid sequence of the protein ^2^. Although these target signals come in a variety of forms, the targeting sequences for chloroplasts, mitochondria and the secretory organelles are usually located at the N-terminus of a polypeptide chain and cleaved upon entry into the organelle ^3^. Thus, the sequences of these targeting signals once removed have no impact on the function of the mature protein. In addition, there is substantial flexibility in the sequence and length of targeting peptides ^4^ such that a large diversity of sequences can function to target proteins to their intended destination.

From early in the investigation of the protein content of organelles it was noted that many proteins had different isoforms with divergent subcellular localisations. For example, the cytosolic and mitochondrial isoforms of phosphoenolpyruvate carboxykinase proteins in animals ^5^, or the cytosolic and chloroplastic isoforms of sugar phosphate enzymes in plants ^6^. Following the advent of protein, cDNA and genome sequence data it was realised that disparate organellar localisation within protein families could be facilitated by differences in the presence and absence of N-terminal target signals and has in fact been found to occur among many paralogous proteins ^7–11^. In addition to these, a bioinformatic analysis of Arabidopsis gene families identified 239 families that contained two or more members with different predicted subcellular localisations, suggesting that changes in protein targeting may be a common occurrence in evolution ^12^. However, when or how often such changes occur during evolution is unknown.

To address this knowledge gap in plants, prediction of subcellular targeting motifs was carried out for the complete set of proteins from a representative set of 42 plant genomes available in the Phytozome database ^13^. The size of the chloroplast, mitochondrion, secretory and peroxisomal proteome for each species was subsequently inferred (Fig.1, Supplemental File S1). Among angiosperms there was little variation in the proportion of the proteome predicted to be targeted to each subcellular compartment while the early diverging land plants and green algae exhibited more variation. On average in land plants the size of the predicted chloroplast, mitochondrion, secretory and peroxisome proteomes comprised 14% (± 2%), 14% (± 3%), 17% (± 2%) and 0.32% (± 0.05%) of the total proteome, respectively (Fig.1, Supplemental File S1). Predicted proteome sizes are likely to be over- or under-estimates depending on the sensitivity and specificity of TargetP, PredAlgo and PTS prediction (see ^14,15^). Irrespective of this however, these results suggest that the proportion of all proteins that are targeted to organelles has remained stable throughout plant evolution.

**Figure 1.**
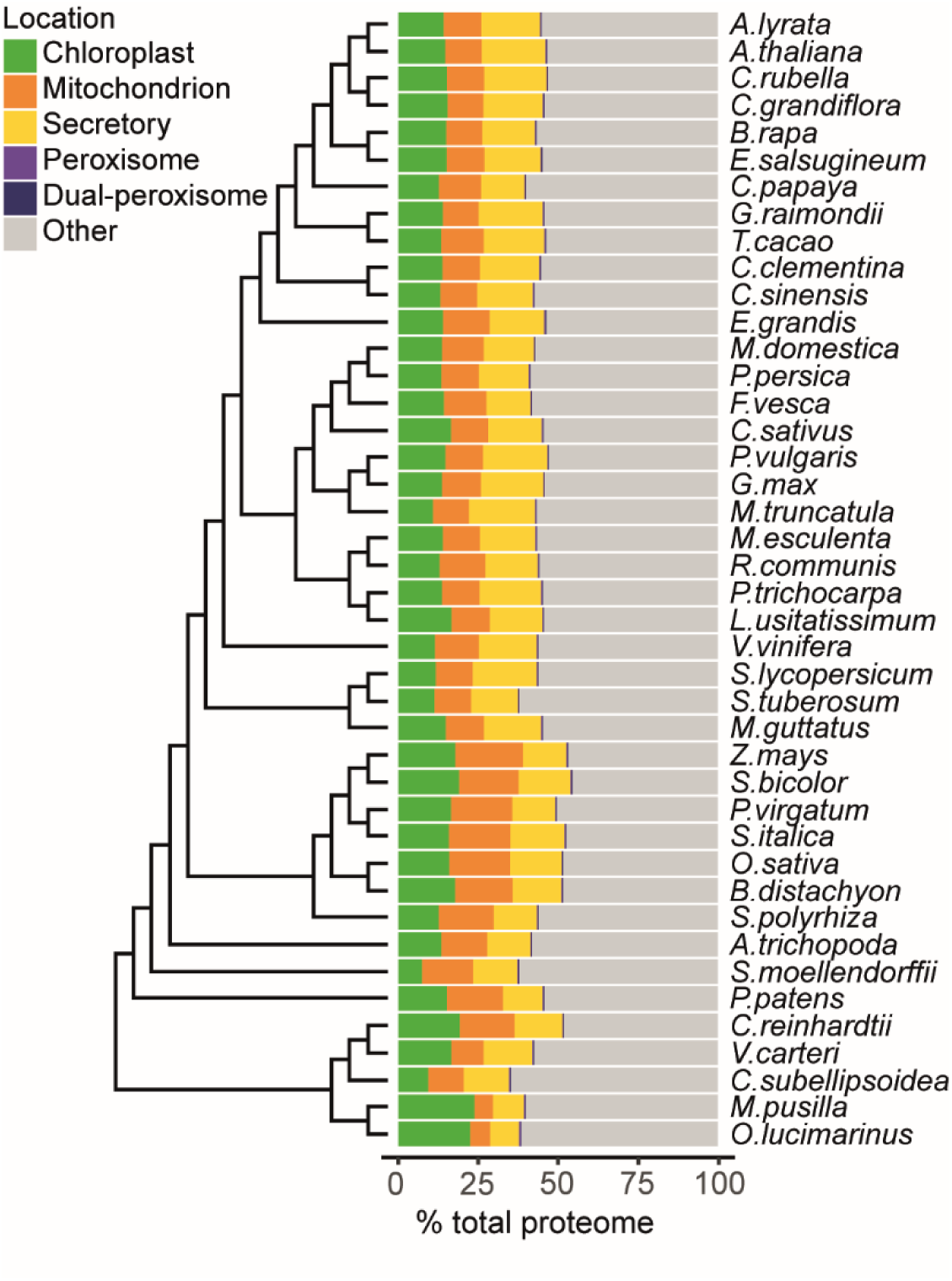
Predicted organelle proteome sizes for each species given as a percentage of the total proteome size of that species. Proteins with both a peroxisomal targeting signal and another predicted target signal (TargetP) were assigned as dual-localised peroxisomal proteins (n = 2973).

To identify occurrences of protein target signal gain and loss during the evolution of plants we inferred a complete set of species-tree reconciled gene trees (n = 18,823) for all orthogroups (gene families) of this 42 species dataset. Ancestral state estimation was then performed to predict the subcellular targeting of the ancestral proteins represented by each internal branch of each reconciled gene tree. Evolutionary changes in protein targeting were identified in this data and mapped to the corresponding branch of the species tree to infer the number of protein gains and losses that occurred to each organelle along each branch of the species tree. In total, across the four organelles, 6162 gains and 9058 losses were identified and mapped to internal branches of the species tree. Gains and losses in protein targeting were observed along every branch of the species tree, with some branches being associated with more change than others (Fig. 2). Incorrect or missing prediction of organelle proteins are a potential source of error in this analysis. To account for this, only changes in protein localisation which have been retained in a high percentage of descendant proteins were selected for this and further analysis (see Methods). This filtration step was included to reduce false positive inference of subcellular localisation change, at the expense of missing some true changes in protein localisation.

**Figure 2.**
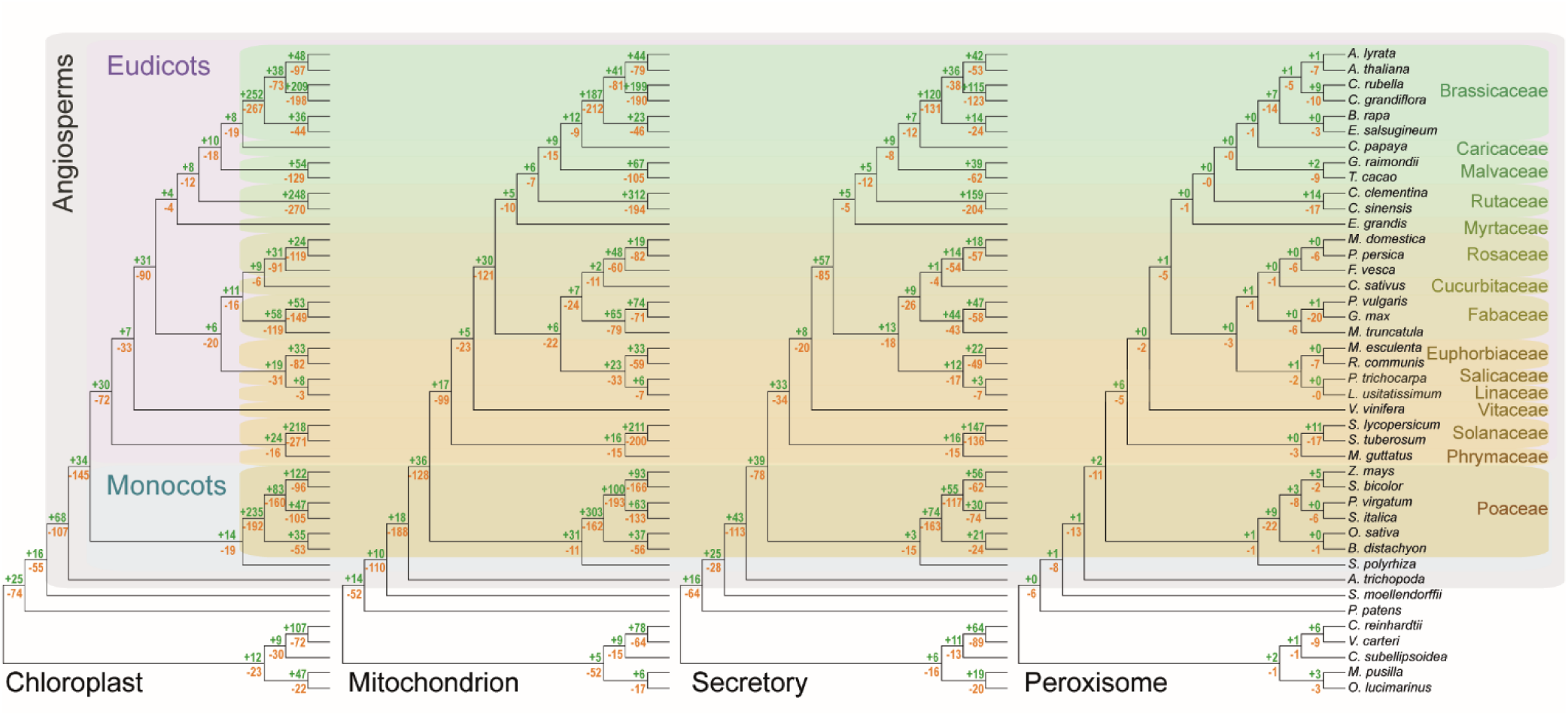
The number of gains (green) and losses (orange) in protein targeting to the chloroplast, mitochondrion, secretory pathway, and peroxisome along each nonterminal branch during the evolution of the species in the study. Note that branch lengths do not correspond to evolutionary distance.

To investigate the rate at which gains and losses in subcellular targeting have occurred during plant evolution the number of changes in subcellular targeting along each branch of the species tree was compared to the amount of molecular evolution that occurred along the same branch. Here the amino acid substitution rate per site was taken as a proxy for molecular evolution rate. There was a positive linear correlation between sequence evolutionary rate and the number of changes in localisation to all subcellular compartments (Fig. 3a-d). Thus, the rate of subcellular targeting evolution is proportional to the rate of molecular evolution and therefore organellar protein content diversifies in proportion to evolutionary distance.

**Figure 3.**
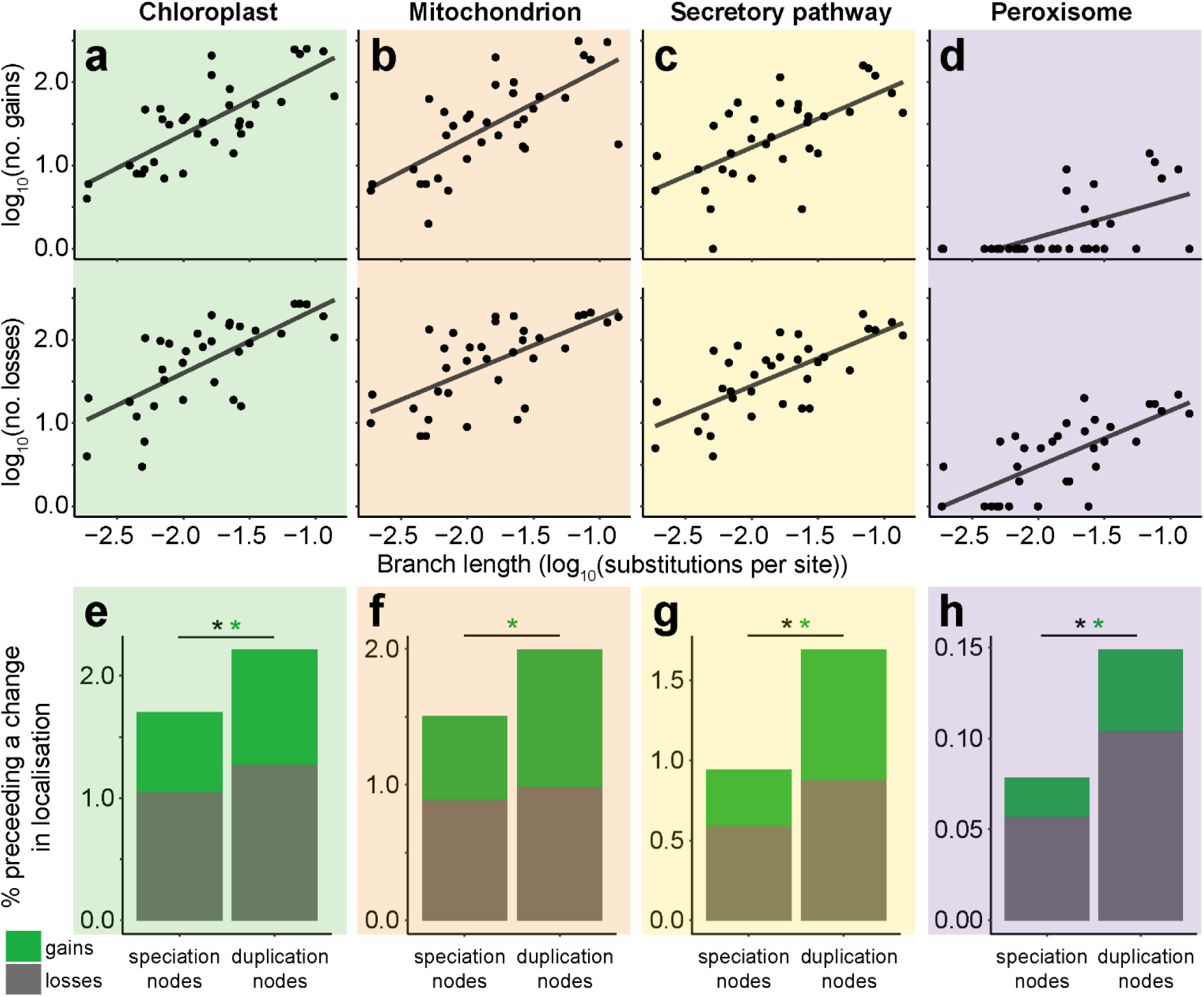
The relationship between evolutionary rate and organellar proteome evolution. There was a positive relationship between species tree branch length (amino acid substitutions per site) and the number of gains or losses in **a**) the chloroplast (R^2^ = 0.59, 0.49), **b**) the mitochondrion (R^2^ = 0.50, 0.42), **c**) the secretory pathway (R^2^ = 0.40, 0.50). All correlations p < 0.001. **d**) fewer gains and losses were observed in peroxisomal targeting, with some branches being associated with no peroxisomal changes, the data is shown but no statistical conclusions drawn. The difference in rates of change in organellar targeting following speciation or gene duplication events in **e**) the chloroplast, **f**) the mitochondrion, **g**) the secretory pathway, **h**) the peroxisome. * indicates significant difference p < 0.01.

While the number of gains along the branches of the species tree was correlated with the number of losses, there was a higher rate of loss in subcellular targeting to each of the four organelles during the evolution of the species in this study (Supplemental Figure, S1). A similar phenomenon was also observed for the gains and losses of signal peptides during the evolution of *Enterobacterales* ^16^. This observation is compatible with the general genetic phenomenon that it is easier to evolve loss-of-function than gain-of-function and thus mirrors studies that have looked at gene or trait gain and loss. Assuming plants colonised the land ~450 million years ago we can estimate that, at a minimum, 3.2, 3.3, 2.2 and 0.21 changes in protein targeting to the chloroplast, mitochondrion, secretory pathway and peroxisome occur for every million years of land plant evolution, respectively (Supplemental File S2). To shed light on the functional significance of these changes in protein targeting, a functional term enrichment analysis was conducted on the set of genes whose localisation changed during plant evolution. For both the chloroplast and the mitochondrion the set of genes that changed localisation during evolution (when compared to the complete set of proteins predicted to be localised to that organelle) were found to be enriched for functional terms concerning regulation, both at a transcriptional and post-transcriptional level (Fig. 4). There was also an overrepresentation of functional terms concerning hormone production, secondary metabolism, stress, transport and development (Supplemental File S3), with few terms related to energy metabolism. In support of this observation, among proteins gained and lost to the chloroplast there was also an over-representation of proteins that localise to the nucleoid, with no statistical over-representation of proteins that locales to other chloroplast sub-compartments such as thylakoid, envelope, or stroma (Supplemental File S4). Thus, it appears that altered regulatory capacity has been the most frequent target of change during the evolution of chloroplasts and mitochondria in land plants.

**Figure 4.**
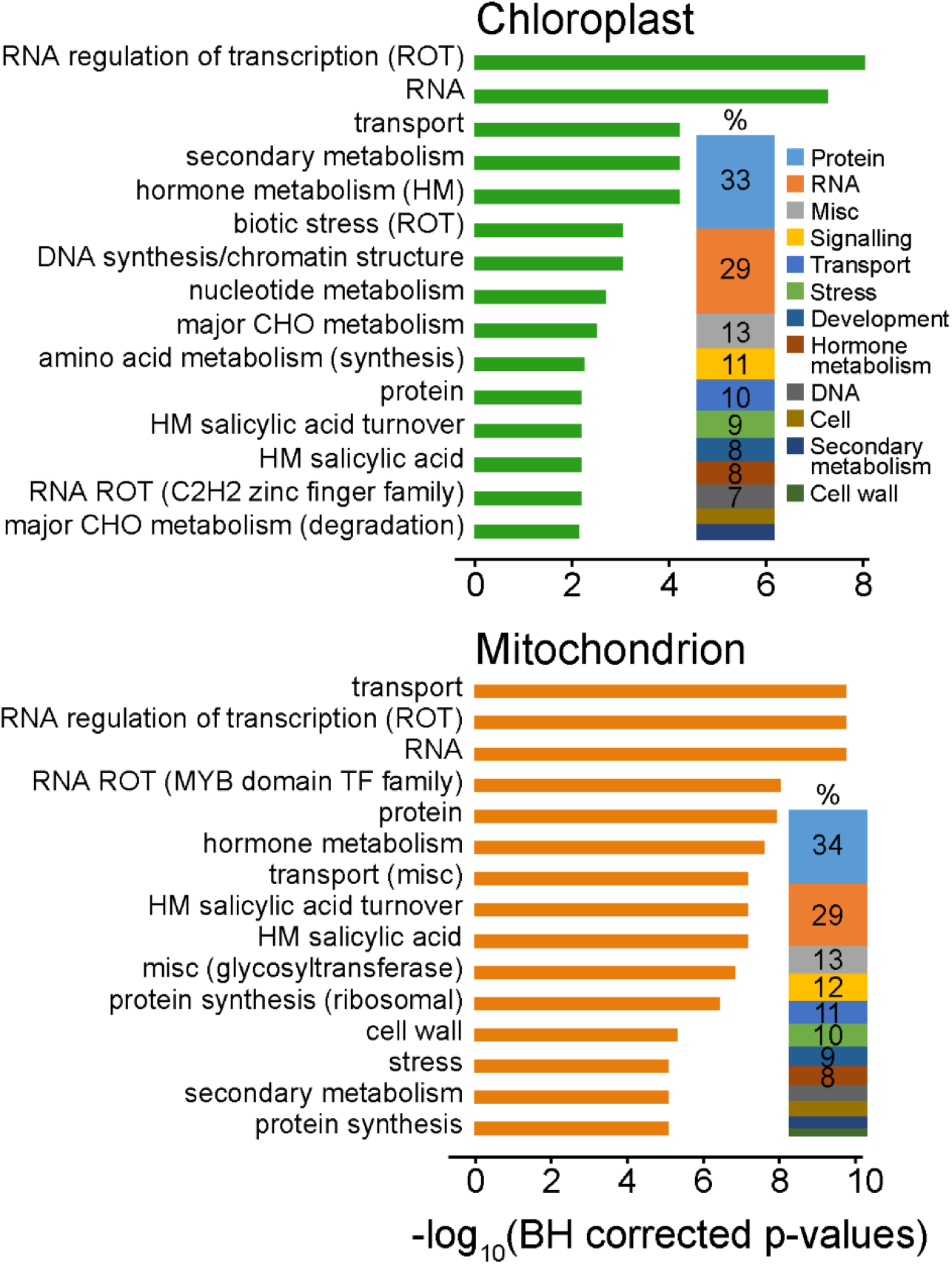
Enriched functional terms (GOMapMan) for the set of proteins that gained or lost a chloroplast or mitochondrial transit peptides during the evolution of the 42 plant species. The top 15 terms are shown for display purposes and the full dataset is available in Supplemental File S3. The proportion plot next to the bar plot indicates the percentage representation of top level functional categories encompassed by the full set of enriched functional terms.

Consistent with the lack of genetic material, functional terms associated with transcriptional regulatory processes were not observed for either the peroxisome or secretory pathway (Supplemental File S3). Instead, enriched functional terms for peroxisomal proteins were associated with metabolism (amino acid, lipid, secondary) or gluconeogenesis while the secretory system were associated with protein post-translational modification, signalling and the cell wall (Supplemental File S3). It was noteworthy that there were a larger number of enriched functional terms for proteins gained or lost to the secretory pathway than any other organelle, consequently there was also a higher diversity of functional classes of genes compared to those relocalised to the chloroplast or mitochondrion (Supplemental File S3).

It has been previously hypothesized that changes in protein localization following gene duplication may be an important mechanism of duplicate gene neofunctionalisation ^7,17–19^. If these hypotheses are correct, it might be expected that changes in protein targeting would evolve more frequently following gene duplication events. To test this, the association between gene duplication events and protein relocalisation events in this data-set was investigated (see Methods). The 18,823 orthogroup trees in this study were analyzed to identify highly-supported, non-terminal gene duplication and speciation nodes. In total 20,137 such duplication nodes were identified and of those 1117 (5.6%) had a child node on which the localization of the protein changed. This frequency was significantly higher than that observed for speciation nodes in the same gene trees (3.9%, p < 0.01). This phenomenon is observed whether the dataset is analyzed as a whole or whether individual locations are analyzed individually (Fig. 3e-h). The one exception to this was the loss of protein targeting to the mitochondrion, which was not significantly higher following gene duplication (Fig. 3f, p = 0.11). Thus, overall the frequency of evolving a change in subcellular localization is higher following gene duplication suggesting that gene and genome duplication may accelerate the pace of organelle proteome evolution.

This study has provided new insight into the dynamics of organellar proteome evolution in plants. It has demonstrated that there has been continuous change in predicted organellar proteomes since plants colonized the land ~450 million years ago. Furthermore, it has revealed that the evolutionary history of the chloroplast and mitochondrion in land plants has primarily been a story of altered regulatory capacity, with the majority of changes occurring to proteins with post-translational or post-transcriptional regulatory functions. The study revealed that the change in organellar proteome content is proportional to the rate of molecular sequence evolution such that plants have gained or lost ~3.2 proteins per million for both the chloroplast and mitochondrion. Finally the study provides evidence that gene duplication leads to enhanced rates of gain and loss of organellar targeting revealing a key role for these events in the evolution of plant organelles.

## Supplemental File legends

**Supplemental Figure S1.** PDF. The ratio of gains to losses for each organelle for each branch in the species tree. Probability density functions were inferred using the density function in R.

**Supplementary File S1.** Microsoft Excel spreadsheet. Sheet 1 (Proteome Sizes) contains the number of genes that encode proteins predicted to be targeted to each subcellular compartment for each species. Sheet 2 (Land Plants Only) contains only data for land plants.

**Supplementary File S2.** Microsoft Excel spreadsheet. Estimation of time calibrated rate of gain and loss. Sheet 1 (Gains and losses (species tree)) contains the number of gains and losses mapped to each node in the species tree for each subcellular compartment. Sheet 2 (Divergence times) contains the divergence time estimates and the number of changes that occurred since that time.

**Supplementary File S3.** Microsoft Excel spreadsheet. Enrichment testing results. There are separate sheets for gains, losses and both combined for each of the subcellular compartments. There are also summary sheets (*_top_level_terms) that contain all of the top level terms for all significantly enriched MAPMAN categories in each *_relocalisations sheets. These summary sheets provide the data for the bar plot in main text Figure 4.

**Supplementary File S4.** Microsoft Excel spreadsheet. Plastid Proteome database Ontology term analysis. Sheet 1 (Ontology_terms) contains a hierarchical representation of the ontology terms provided in the plastid proteome database. Sheet 2 (PPDB_data) contains all of the PPDB data downloaded on the 13^th^ of March 2018, it is provided here for reference in the event that the database is lost or updated. Sheet 3 (Orthogroup_PPDB_terms) contains the ontology term to orthogroup mapping used in this analysis. Orthogroups inherit an ontology term if they contain a gene which has that ontology term.

## Acknowledgements

RC is supported by a BBSRC studentship through BB/J014427/1. SK is a Royal Society University Research Fellow. Work in SK’s lab is supported by the European Union’s Horizon 2020 research and innovation program under grant agreement number 637765.

## Online Methods

### Data availability

All data used in this study has been deposited, and is freely available, at the Zenodo research data archive https://doi.org/10.5281/zenodo.1414180. This archive contains the full set of sequences, accession numbers, predicted localisation data, orthogroups, and PHYLDOG reconciled gene trees for each orthogroup. The archive also contains the full information for all of the gene duplication events and change in localisation events for each orthogroup.

### Proteome data, and inference of orthogroups and gene trees

The protein primary transcripts for 42 fully sequenced plant species were obtained from Phytozome v10^13^. Orthologous gene groups (orthogroups) were inferred using OrthoFinder ^20^ and multiple sequence alignments were inferred for each orthogroup using MAFFT-LINSI ^21^. To minimise the contribution of gene tree inference error, gene trees were inferred using the true species tree for guidance by simultaneous gene tree-species tree reconciliation using PHYLDOG ^22^. PHYLDOG was run on each orthogroup alignment individually with a user-provided species tree. The “LG08” model was used, the maximum number of gaps allowed in an alignment column was 66%. The topology of the species tree was derived from angiosperm phylogeny working group and branch lengths were inferred from a concatenated multiple sequence alignment with the topology constrained to the this topology tree using RAxML ^23^. All attempts to jointly infer the gene trees and species tree by analysing all the orthogroupstogether resulted in an error (“Assertion ‘tr->likelihood >= currentLikelihood’ failed.”), hence all orthogroups were analysed individually with the species tree as input. The largest orthogroups could not be analysed directly with PHYLDOG as they were too large (the largest orthogroup contained 12148 genes). To allow the whole dataset to be analysed, gene trees for the largest 100 orthogroups were inferred using FastTree ^24^. The nodes of these gene trees were mapped to the species tree using DLCpar ^25^. The orthogroups were then split at each node corresponding to the root of the species tree, and each of these splits were analysed separately.

### Prediction of organelle proteins

Of the 42 species included in this study, 37 comprise land plants and five comprise green algae. From proteome data for each species, we looked to identify the set of proteins predicted to contain a chloroplast transit peptide (cTP), mitochondrial targeting peptide (mTP), signal peptide (SP) or the peroxisomal targeting signals 1 & 2 (PTS1 & PTS2). For the land plant species cTPs, mTPs and SPs were predicted by TargetP 1.1 ^14^ in plant mode with default cutoffs. For the five algal species (*O. lucimarinus, M. pusilla, C. subellipsoidea, V. carteri, C. reinhardtii*) this prediction was carried out with PredAlgo ^15^ using its default cutoffs. In cases where an amino acid sequence did not meet the minimum length requirement for PredAlgo prediction, the TargetP prediction was taken instead.

The prediction of peroxisomal proteins was carried out by searching for the canonical plant peroxisomal targeting signals 1 and 2 (PTS1 and PTS2) ^26^. Here a protein sequence was classified as having a PTS1 if it had any one of the 9 different c-terminal tripeptide sequences (SRL, SRM, SRI, ARL, ARM, PRL, SKL, SKM, AKL). Similarly, a protein sequence was classified as having a PTS2 peroxisome targeting sequence if it contained either of the two PTS2 peptide sequences (R[LI]X5HL) in the N-terminus region of the protein (residues 1 – 30).

### Ancestral character estimation of subcellular targeting

Maximum-likelihood ancestral character estimation was used to identify gain and loss events in protein targeting that occurred during the evolution of the orthogroups inferred from this dataset. Considering the four types of target signal separately, the presence or absence of a predicted target sequence within each protein was treated as binary trait data and the leaf nodes of orthogroup trees assigned a “1” or “0” accordingly. Ancestral character estimation was then carried out independently for each orthogroup to estimate the character state (presence/absence of a targeting sequence) of each internal node using the “ace” function in the R package ape ^27^ for discrete data and using the “all rates different” model. The model selected for ace is dependent upon the transition probabilities between the states. For binary characters either an “equal rates” model, in which the transition between states is constrained to be equal, or an “all rates different” model in which the forward and backward transition rate was allowed to vary over the tree can be used. It is unknown whether the rate at which a protein can gain or lose a target signal is equal, therefore the “all rates different” model was selected as being most appropriate for ancestral state estimation. Internal nodes in orthogroup trees with likelihood scores ≥0.5 were considered to contain a targeting sequence, while nodes with scores <0.5 were considered to lack a targeting sequence. Further processing and filtration was carried out as described below.

### Identifying changes in the subcellular localisation of a protein during evolution

The ancestral character estimation data was analyzed to identify changes in the organellar targeting of proteins in each orthogroup tree. By iterating over the internal nodes of the tree, a loss in subcellular targeting was defined as a transition from a targeted state to not-targeted state on consecutive branches, and *vice versa* for a gain. As ancestral character estimation is sensitive to prediction or gene tree error, a stringent filter was imposed such that a change in subcellular targeting was only counted if the changed state was conserved in 75% of the genes below the node on which the change occurred. For example, consider a parent node X and two child nodes Y and Z. If there was a predicted gain of a chloroplast transit peptide between node X and one of its child nodes Y, then 75% of the proteins on the branches that subtend node Y must contain a predicted chloroplast transit peptide for it to be considered for further analysis. Similarly, for the other child node Z, 75% of the genes that subtend that must not contain a chloroplast transit peptide. Only if both these criteria are met is a change in subcellular localisation assigned to the branch within the orthogroup tree between node X and node Y. In all cases, it was required that two or more sequences must subtend any branch under consideration.

This requirement was imposed so that inference about the predicted subcellular targeting state of an ancestral protein was informed by the subcellular targeting state of two or more extant genes. This requirement improves robustness to subcellular prediction targeting error and means that changes in subcellular localisation was not evaluated for terminal branches in orthogroup trees. This filtered dataset was used in all subsequent analyses.

Given that each orthogroup tree was reconciled with the species tree, the complete set of changes in all orthogroup trees could be assigned to the corresponding branches on the species tree. Thus the number of gains and losses in protein targeting to each of the four organelles could be quantified for each branch of the species tree.

To estimate the average rate at which proteins have gained or lost organelle target signals during the evolution of land plants 10 nodes were selected on the tree for which a divergence time is known. The number of gains and losses in targeting to each organelle was then summed for the branches between the node at the base of the land plants (taken as 450mya ^28^) and the nodes with known divergence time, thus allowing the number of changes per million years to be calculated (Supplementary File S2).

### Identification of changes following gene duplication and speciation events

To investigate whether changes in subcellular targeting occur more frequently following gene duplication events or speciation events it was necessary to identify whether each node in each orthogroup tree comprised either a gene duplication event or a speciation event. To prevent tree inference error from influencing the results, a stringent filter was applied to the orthogroup trees to enable identification of both high confidence gene duplication nodes and high confidence speciation nodes. High confidence gene duplication nodes were defined as nodes for which the gene duplication event was retained in all descendant species of both child nodes subtending the gene duplication event. Similarly a high confidence speciation node was selected as a node which has no evidence for gene duplication and from which there was no subsequent gene loss in any of the descendant species. In both cases, (duplication and speciation nodes) complete retention of all genes in all descendant species is required and thus the gene sets can be considered broadly equivalent.

To determine whether changes in subcellular localisation were more likely to occur following gene duplication events than speciation events, the occurrence of changes in subcellular localisation following duplication or speciation nodes was analyzed.

### Functional term enrichment analysis

Orthogroups were assigned MapMan terms and sub-chloroplast localisation terms (plant protein database PPDB) by inheriting the terms associated with the genes found within them. MapMan terms were taken from the GoMapMan webpage ^29^ and sub-chloroplast terms assigned using the hierarchical structure provided on the PPDB ^30^ using only experimentally validated proteins (see Supplementary File S4 for the PPDB list used at time of writing). To test for enrichment the hypergeometric test was performed and p-values corrected for multiple testing using the Benjamini-Hochberg correction (see Supplementary File S3 for MapMan results and S4 for PPDB). The aim was to identify functional enrichment among orthogroups whose proteins are differentially localized. To avoid simply identifying functional terms that are enriched in organelle targeted gene families, the background sample for this test was orthogroups with at least one predicted organelle targeted protein. Significantly enriched functional annotation terms were those with a corrected p-value of ≤ 0.01.

